# Sex-Related Associations Between Precuneus Volume and Subjective–Objective Memory Discrepancy in Amyloid-Positive Individuals

**DOI:** 10.64898/2026.11.11.766001

**Authors:** Marc-Antoine Akzam-Ouellette, Vladimir Fonov, Reza Rajabli, Madeleine Sharp, D. Louis Collins, the Alzheimer’s Disease Neuroimaging Initiative

## Abstract

**INTRODUCTION:** The precuneus, a metamemory-related parietal region with sex-specific vulnerability in Alzheimer’s disease (AD), may contribute to reduced insight of memory deficit. We investigated whether precuneus structural integrity explained sex differences in subjective-objective memory discrepancy.

**METHODS:** Precuneus volumes, as well as three other control regions, were acquired in 350 Aβ+ and 310 Aβ-participants from the ADNI dataset to quantify within-individual changes in subjective–objective memory discrepancy across AD progression.

**RESULTS:** Aβ+ participants exhibited steeper objective memory decline (*β* = −0.446, *p* < 0.001) despite stable subjective complaints (*β* = −0.027, *p* = 0.587), resulting in a widening subjective-objective memory discrepancy (*β* = 0.317, *p* < 0.001). In this group, only precuneus atrophy showed a sex-dependent association with discrepancy (*β* = 0.508, *p* = 0.002), a finding absent in the Aβ− group or other brain regions.

**DISCUSSION:** The precuneus represents a neuroanatomical substrate associated with sex-related differences in metacognitive vulnerability in Aβ+ individuals.

## Introduction

By 2050, over 152 million people worldwide are expected to be affected by dementia, with Alzheimer’s disease (AD) as the most prevalent (Guerchet et al., 2020). Growing evidence identifies that after age, sex is a key factor in AD progression. Yet, sex-specific trajectories and their cognitive and behavioural phenotype remain to be fully defined (Lopez-Lee et al., 2024). This need for clarification is exemplified in AD: females APOE4 carriers have higher CSF t-Tau and p-Tau concentrations in early stages than males (Babapour Mofrad et al., 2020), and faster cognitive decline after a diagnosis of Mild Cognitive Impairment (MCI) (Ferretti et al., 2018). An understudied aspect of AD’s clinical profile regarding sex differences is subjective cognitive complaints (SCC), an individual’s perceived cognitive decline that serves as a key diagnostic criterion for future amnestic MCI, the prodromal stage of AD (Petersen & Morris, 2005). SCC is also associated with an increased likelihood of preclinical AD (Jessen et al., 2020), where overestimation of memory abilities correlates directly with the severity of cognitive decline in MCI (Perrotin et al., 2007). We hypothesize that a form of anosognosia (i.e., a lack of awareness or insight into one’s own deficits) may be present in some subjects on the AD trajectory, potentially causing a discrepancy between subjective and objective measures of cognitive deficit (Buckley et al., 2015).

Sex could influence this discrepancy and therefore might impact the validity and utility of SCC. Analyzing 940 controls and aMCI participants from the Alzheimer’s Disease Neuroimaging Initiative (ADNI), Sundermann et al. (2018) found that sex moderated the predictive validity of memory complaints at the aMCI stage. They employed a discrepancy score that compares the participant’s self-report to an informant’s assessment on the Everyday cognition scale (ECOG) (Farias et al., 2008) to capture the gap between self-perceived and informant-perceived cognitive abilities. In aMCI females, self-reported complaints showed a strong relationship with objective memory performance. For aMCI males, the relationship was weaker, with a tendency to underestimate their memory deficits. These results are in line with those of Coddaire et al. (2025), who found a similar pattern in MCI participants with the same methodology, but they did not examine sex-specific effects.

A key region that could be implicated in memory deficit insight is the precuneus, a parietal structure within the default mode network (DMN) associated with metamemory processes (Park et al., 2018). The precuneus plays a central role in self-referential processing and efficient retrieval of episodic memories (Cavanna & Trimble, 2006; Ren et al., 2018)) and has been shown to have a causal role in mnemonic metacognition (Ye et al., 2018). This region is affected early in AD, with DMN–precuneus connectivity specifically disrupted (Klaassens et al., 2017; Wang et al., 2024), and is among the first areas to show Aβ accumulation (Ali et al., 2022; Palmqvist et al., 2017), and impaired functional within-network connectivity in this region is associated with anosognosia in MCI (Mondragón et al., 2019).

Several studies also suggest sex differences in the role played by the precuneus. Zhang & Li (2012) found that females show greater ventral precuneus-hippocampal connectivity than males. Building on this, Cieri et al. (2022) showed that females tend to maintain cortical thickness in DMN regions, including the precuneus, from the preclinical to the prodromal stage, but then undergo a steeper decline from prodromal stage to dementia. Whether sex differences in the vulnerability of the precuneus to AD could contribute to sex differences in the insight of memory differences remains unknown.

As these discrepancies may evolve with AD progression, the objectives of this study were to use a longitudinal approach to examine whether baseline precuneus integrity and subsequent atrophy differentially predict the alignment between subjective complaints and objective memory performance between sexes. We hypothesized that females maintaining greater precuneus structural integrity would show stronger alignment between subjective and objective memory than males, both cross-sectionally and longitudinally.

## Methods

### Participants

Data used in the preparation of this article were obtained from the Alzheimer’s Disease Neuroimaging Initiative (ADNI) database (adni.loni.usc.edu). The ADNI was launched in 2003 as a public-private partnership, led by Principal Investigator Michael W. Weiner, MD. The primary goal of ADNI has been to test whether serial magnetic resonance imaging (MRI), positron emission tomography (PET), other biological markers, and clinical and neuropsychological assessment can be combined to measure the progression of mild cognitive impairment (MCI) and early Alzheimer’s disease (AD) (Mueller et al., 2005). For up-to-date information, see www.adni-info.org.

Participants were drawn from ADNI-1, ADNI-2, ADNI-3 and ADNI-GO as part of a longitudinal design and separated in two groups according to their phospholaryzed-tau (pTau)/Aβ(1-42) level status ratio when both biomarker measures were available. A participant with a pTau/Aβ ratio above 0.028 at any given time point was considered Aβ positive (Aβ+) and on the AD continuum (Jack Jr et al., 2024). This ratio is based on cut-offs derived from BioFINDER and adjusted for pre-analytical handling in ADNI (Hansson et al., 2018). A participant that did not reach that ratio at any time point was considered Aβ negative (Aβ−; below or equal to 0.028) and grouped as such. Participants were stratified by amyloid status to distinguish AD–related pathology from non-AD causes of cognitive impairment. This allowed us to examine the effect of AD-specific pathology on precuneus structural integrity while minimizing confounding from other dementias (i.e., frontemporal dementia, vascular dementia, etc.).

To examine the relationship between precuneus integrity and subjective–objective memory discrepancy across the AD trajectory, we used complementary baseline and longitudinal approaches. Baseline analyses assessed whether initial precuneus integrity was associated with memory insight and whether these associations differed by sex, whereas longitudinal models examined within-individual changes over time and whether progressive structural decline differentially influenced this relationship between sexes. Participants in the Aβ+ group had on average 2.5 visits (SD = 1.59), and participants in the Aβ− group had an average of 3.3 visits (SD = 1.87). Cognitive metrics were z-scored using the mean and standard deviation of the cognitively unimpaired Aβ− participants (recall that ADNI also has Aβ− MCI and Dementia participants) at their first available assessment. This standardization provided a normative reference, allowing scores to be interpreted as deviations from a cognitively healthy, amyloid biomarker-negative sample.

The primary inclusion criterion was the availability of both pTau and Aβ(1−42) measurements at one or more study visits, allowing calculation of the pTau/Aβ(1−42) ratio used for group classification; as well as at least two visits with MRI measurements. No additional inclusion or exclusion criteria were applied. Standard ADNI cohort descriptions and criteria are detailed at https://adni.loni.usc.edu/data-samples/adni-data/study-cohort-information/. All ADNI participants provided informed consent in compliance with the Declaration of Helsinki, and ethical approval was obtained from the institutional review boards of each participating site.

### Memory Assessments

#### Discrepancy

While the aforementioned articles (Coddaire et al., 2025; Sundermann et al., 2018) provided valuable insights into SCC, their methodology for calculating a subjective−objective discrepancy metric presents a crucial limitation. Their scores were derived from the difference between the participant’s self-report and the informant’s assessment. This approach captures the difference in perception between two individuals’ subjective evaluation rather than the essential divergence between a participant’s *subjective* experience and their actual, *objectively* measured, memory performance. This reliance on an informant-based measure poses two challenges for investigating neural correlates, such as the precuneus: it risks incorporating external informant subjective bias and does not isolate core memory performance integrity. Calculating the discrepancy by subtracting the participant’s objective assessment of their memory ability from a subjective memory measure may provide a more accurate estimate of cognitive discrepancy, reflecting the gap between perceived and actual memory performance. To evaluate self-assessment of memory performance, a *cognitive* discrepancy score was created and calculated, based on the difference between a subjective memory measurement and an objective metric, further detailed below.

To account for differences in disease severity at study entry, baseline clinical status was indexed by the Clinical Dementia Rating–Sum of Boxes (CDR-SB) (Morris, 1993). Including baseline CDR-SB as a covariate accounted for variability in participants’ clinical stage at their individual baseline, allowing longitudinal trajectories to be estimated independently of initial cognitive impairment.

#### Subjective Memory

To assess subjective memory performance, items of the different memory subscales of the ECOG were used, as done in the two previously mentioned studies. The ECOG is a validated subjective measurement of cognition, based out of 39 items that target various domains, such as memory, orientation, and planification (e.g., for memory: *Remembering a few shopping items without a list*; *Remembering appointments, meetings, or engagements*, etc.). All items are based on a 4-point scale, going from 1 to 4. A score of 1 indicates *better or no change* for that specific item, a score of 2 indicates *questionable/occasionally worse*, a score of 3 indicates *consistently a little worse*, and a score of 4 indicates *consistently much worse*. For the subjective measure, we used a subset of 8 ECOG memory-related items previously shown to discriminate between cognitively normal participants and individuals with MCI (Farias et al., 2008). The total sum of these 8 memory items then ranges from possible values between 8 (unimpaired) and 32 (severely impaired). This sum served as the subjective metric, which is reversed so that large positive values indicate better subjective performance. This subjective score was standardized (z-scored) against the results of the Aβ− reference sample.

#### Objective Memory

To assess objective episodic memory performance, a composite score based on the average of the Weschler logical memory immediate and delayed recall was computed (Wechsler, 1987). The reference sample comprised those cognitively unimpaired Aβ− participants (based on ADNI diagnosis and the pTau/Aβ(1–42) ratio) that had valid immediate and delayed recall and ECOG scores at their first available assessment, ensuring all measures required to compute the memory–subjective cognition discrepancy score (detailed below) were available.

Both the immediate and the delayed recall were standardized (z-scored) against the performance of the Aβ− reference sample. The standardized delayed recall score and the standardized immediate recall score were averaged at each time point across participants to provide the objective memory metric. Again, larger positive values indicate better performance.

#### Cognitive Discrepancy Score

To quantify the misalignment between subjective and objective memory performance, a *cognitive discrepancy* score was computed. This score was derived by subtracting the standardized objective memory metric from the standardized subjective memory rating. Figure 1 show the calculation steps for the Discrepancy score for both groups.

**Figure 1.**
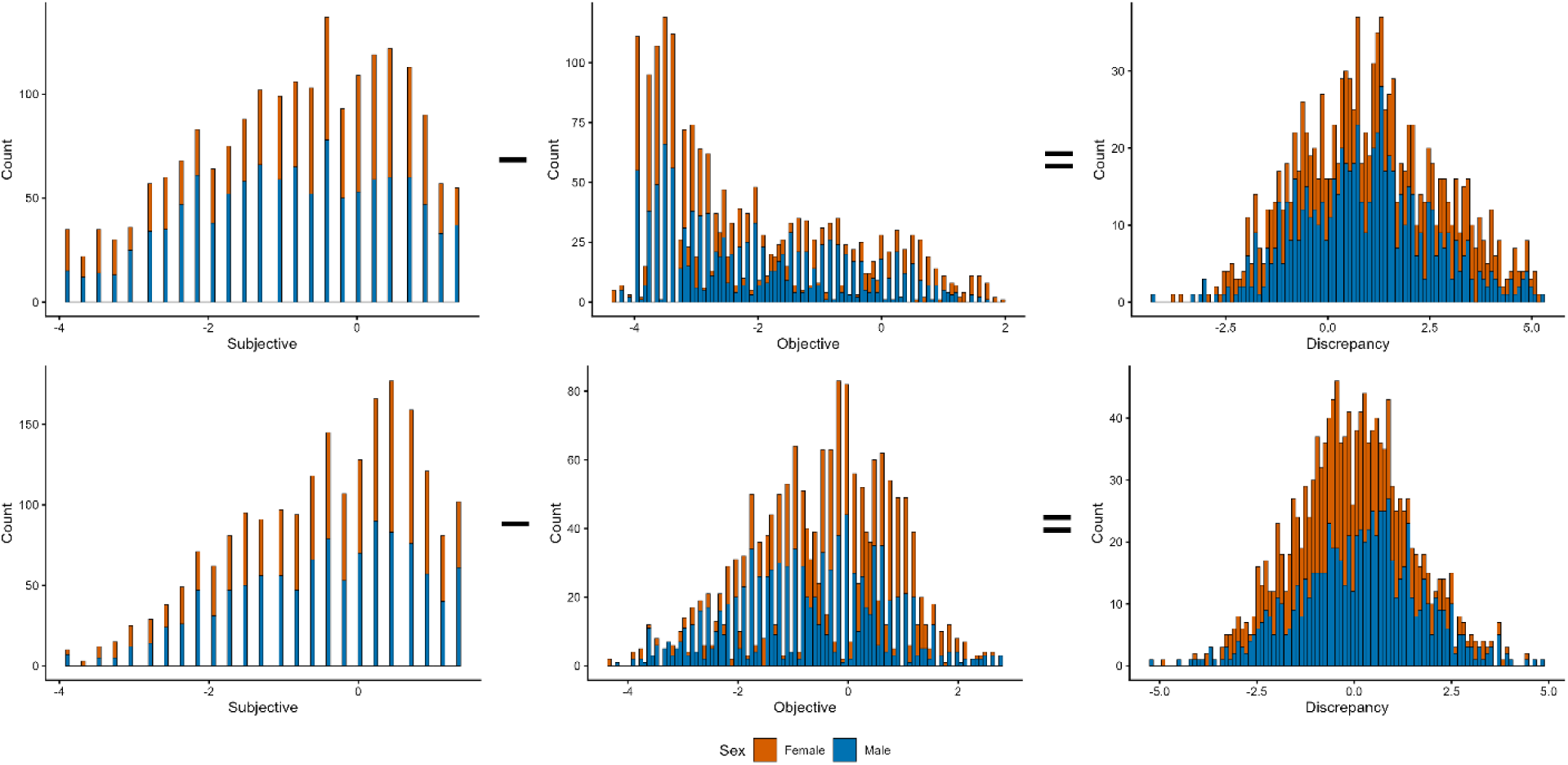
Discrepancy Calculation by Subtracting Objective from Subjective performance in Each Group. Note. First row are results for the Aβ+ group. Second row is for the Aβ− group.

### Imaging

All ADNI participants received a T1-weigthed (T1w) MRI scan at each time point (see http://adni.loni.usc.edu/methods/mri-tool/mri-analysis/ for the complete acquisition protocol). All scans were inspected prior to preprocessing for major artifacts. Brain regions were segmented using AssemblyNET (Coupé et al., 2020), a validated deep-learning segmentation framework. Registration quality control was performed automatically using RegQCNet, a deep convolutional neural network that quantifies affine registration error between each registered image and the reference template in metric units (Denis de Senneville et al., 2020). Based on this quantitative error estimate, scans were categorized into quality tiers, and only those rated as excellent were included in subsequent analyses. While we are interested in the precuneus, three additional brain region volumes were included to serve as controls. The first control brain region is the hippocampus (HC), since this region has been shown to the affected in AD, and is critically involved in objective memory impairment. The second control region is the superior frontal gyrus (SFG), since it is generally considered not to be affected by AD progression. This choice of control regions is supported by the work of Coupé et al. (2019) and Fonov et al. (2026). The third region is another strong candidate for its potential role in anosognosia in AD, the Posterior cingulate cortex (PCC). Andrade & Pacella (2024) proposed that in AD, anosognosia would arise from a failure in error monitoring triggered by early Aβ deposits in the PCC. These deposits may play a causal role by impairing patients’ insight of their own deficits. This claim is supported by many studies that have shown a relationship with hypometabolism in this region and anosognosia (Perrotin et al., 2015) or that dysregulation of large scale networks and the PCC were associated with greater anosognosia (Tondelli et al., 2024). Thus, if significant results are found with the precuneus, and not with the three other regions, it would be possible to conclude that the former, and not the three latter, is involved in subjective−objective discrepancy between sexes in AD. Intracranial volume (ICV) was also estimated by AssemblyNet and used to control for differences in the total brain size. Raw structural volumes were used, with the left and right summed for each region.

### Data Analysis

Our primary hypothesis was that sex would moderate the association between precuneus integrity and memory insight in Aβ+ individuals, such that greater structural integrity would be more strongly associated with better alignment between subjective and objective memory performance in females than in males. This hypothesis was examined both cross-sectionally and longitudinally, with structure Regional_volume used for a proxy of structural integrity in the cross-sectional model. Change in volume compared to baseline, Regional_delta, was used as a proxy of structural integrity in the longitudinal model (with baseline Regional_volume used as a covariate). Statistical analyses were designed in a stepwise manner to progressively build the final interaction model to contextualize our results according to differences in cognitive performance and brain atrophy trajectories between groups. Prior to model building, baseline demographic and clinical characteristics were compared between Aβ+ and Aβ− groups using independent-samples *t*-tests for continuous variables (e.g., age (years), education (years), baseline CDR-SB score) to identify potential group differences. Time since baseline (as days) was z-scored using the mean and SD of the full analytic sample, prior to stratification by amyloid status, to preserve comparability of the time metric across Aβ+ and Aβ− models.

Linear mixed-effects models, with participant-specific random intercepts and slopes to account for within-participant correlation, characterized differences between Aβ+ and Aβ− individuals at model intercept and over time (Aβ status × Time_z_ interaction). Models of objective memory performance, subjective memory performance, and their discrepancy were adjusted for sex, education, and baseline CDR-SB (eq. 1);

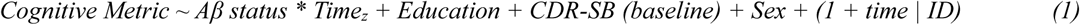

Models of regional brain atrophy across four regions of interest were adjusted for ICV and sex (eq. 2).

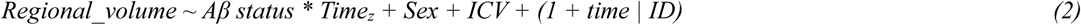

Final analyses tested Regional_volume × Sex interactions on memory insight discrepancy, our outcome measure of interest, separately for the precuneus and for three control regions, cross-sectionally (eq. 3) and longitudinally (eq. 4).

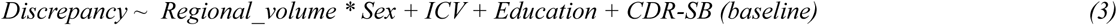

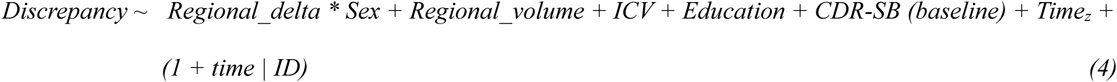

Cross-sectional Discrepancy models (eq. 3) use only baseline data and tested whether Regional_volume × Sex predicted Discrepancy, adjusting for ICV, Education, and baseline CDR-SB, with regional baseline volume mean-centered to reduce multicollinearity. Longitudinal models (eq. 4) tested whether within-subject Regional_delta (i.e., volume at each timepoint minus baseline volume) × Sex predicted change in Discrepancy, adjusting for Time, baseline Regional_volume, ICV, Education, and baseline CDR-SB. Participant-specific random intercepts and slopes accounted for repeated measurements and within-subject correlations.

All discrepancy interaction models (eq. 3 and 4) were conducted separately in Aβ+ and Aβ− individuals to examine the association between regional volume and sex within each group, with a specific focus on the Regional_volume × Sex interaction in Aβ+ individuals, as predicted by our hypothesis.

A *p-*value < 0.05 was considered significant for all analyses. Marginal and conditional R^2^ (Nakagawa & Schielzeth, 2013) were computed to assess effect size of the models. All analyses were done using R statistical software, version 4.2.2. Regression models were conducted using either the *lm* or *lmer* (lme4 package) functions in R (Bates et al., 2015). P-values were calculated with the *lmerTest* package (Kuznetsova et al., 2017).

## Results

### Group Characteristics

The Aβ+ group had 350 participants (154 females), and the Aβ− group had 310 participants (147 females). The reference sample of cognitively unimpaired Aβ− participants was made of 180 participants (91 females). Sex distribution was comparable between amyloid groups (Fisher’s exact test, *p* = 0.390; OR = 1.15, 95% CI [0.83–1.58]). The Aβ+ group was significantly older than the Aβ− group (*t* = −3.235, *p* < 0.01); however, the effect size was small (Cohen’s *d* = −0.25). The Aβ+ group also had lower educational attainment compared with the Aβ− group (*t* = 3.497, *p* < 0.001), although the magnitude of this difference was small (Cohen’s *d* = 0.27). Clinical severity measured by the CDR−SB at baseline was significantly higher in the Aβ+ group compared with the Aβ− group (*t* = −9.742, *p* < 0.001, Cohen’s *d* = −0.65). Table 1 describes the demographic characteristics of both groups and the reference sample at individual baseline, stratified by Sex.

**Table 1.** Demographic Characteristics of the Aβ+, the Aβ− and the Reference Groups at Individual Baseline.

| **Group** | **Sex** | ***n*** | **Age** | **Education** | **Aβ level** | **pTAU** | **CDR-SB<br>(baseline)** |
|---|---|---|---|---|---|---|---|
| **Aβ+** | Female | 154 | 72.76 (6.96) | 15.27 (2.57) | 718.18 (227.02) | 37.99 (16.34) | 1.74 (1.79) |
|  | Male | 196 | 74.45 (6.52) | 16.42 (2.56) | 648.95 (212.61) | 33.22 (12.92) | 1.84 (1.5) |
| **Aβ-** | Female | 147 | 71.11 (7.34) | 15.99 (2.53) | 1184.21 (305.33) | 16.76 (5.65) | 0.65 (0.79) |
|  | Male | 163 | 72.7 (6.89) | 17.18 (2.44) | 1172.79 (325.36) | 16.71 (5.39) | 0.87 (1.23) |
| **Reference** | Female | 91 | 70.70 (6.76) | 16.12 (2.48) | 1273.55 (233.15) | 17.20 (4.03) | 0.21 (0.55) |
|  | Male | 89 | 71.96 (7.00) | 17.29 (2.38) | 1269.44 (247.42) | 18.39 (4.33) | 0.22 (0.52) |

*Note.* Higher CSF Aβ(1-42) values indicate lower brain amyloid burden, as reduced CSF Aβ(1-42) concentrations are associated with increased cerebral amyloid deposition due to sequestration of Aβ(1-42) into plaques (Andreasen et al., 1999). Aβ = amyloid-beta; pTau = phosphorylated tau; CDR-SB = Clinical Dementia Rating Scale–Sum of Boxes. N=number of subjects, age in years, education in years. The reference group is the cognitively unimpaired Aβ− group.

### Memory Performance Across Groups Over Time (eq. 1)

For the objective memory performance, the Aβ+ group exhibited significantly lower scores compared with the Aβ− group (*β* = −1.023, *p* < 0.001). Males had significantly lower scores than females overall (*β* = −0.291, *p* < 0.001). In addition, a significant Aβ status × time interaction was observed (*β* = −0.446, *p* < 0.001) indicating a steeper longitudinal decline in objective memory performance in the Aβ+ group compared with the Aβ− group.

For subjective memory performance, the Aβ+ group showed significantly lower scores compared with the Aβ− group (*β* = −0.316, *p* < 0.001). No Sex differences were found (*β* = −0.123, *p* = 0.145). Furthermore, the Aβ status × Time interaction was not significant (*β* = −0.027, *p* = 0.587), indicating that subjective memory trajectories did not differ between groups over time.

For the cognitive discrepancy interaction model, the Aβ+ group demonstrated significantly higher Discrepancy scores compared with the Aβ− group (*β* = 0.676, *p* < 0.001). No Sex differences were found across groups (*β* = 0.127, *p* = 0.262). A significant Aβ status × time interaction was observed (*β* = 0.317, *p* < 0.001), indicating a greater longitudinal increase in Discrepancy in the Aβ+ group. Figure 2 shows the trajectories of the Subjective, Objective and Discrepancy measurement in time, separated for both amyloid groups. Table 2 shows the results of the different cognitive metrics at baseline for both Aβ+ and Aβ− groups, as well as the reference sample, stratified by Sex.

**Figure 2.**
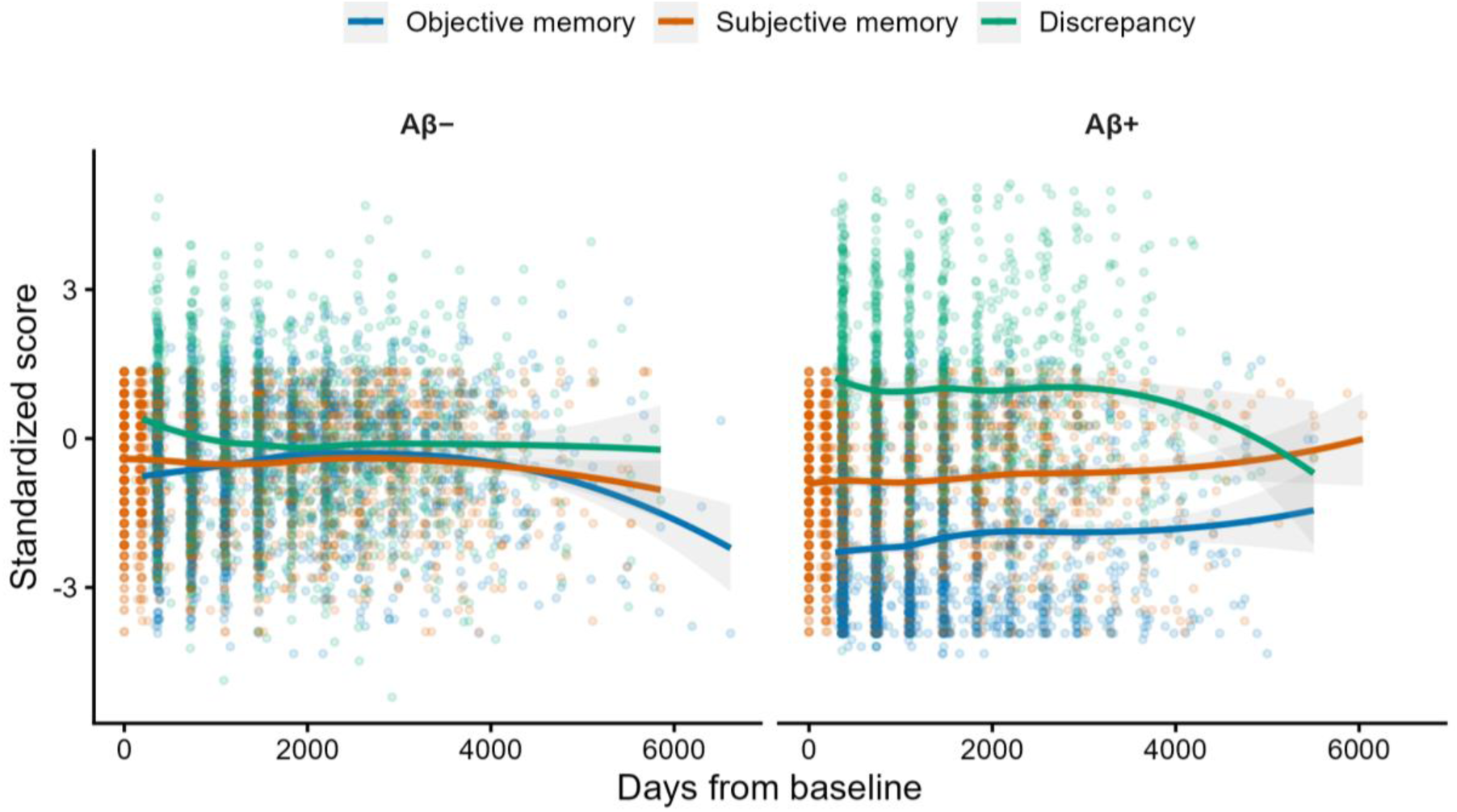
Memory Measures Trajectories in Time Separated for Both Amyloid Groups. *Note.* Objective and subjective memory LOESS trajectories remain aligned in the Aβ− group, yielding discrepancy scores near zero, whereas the Aβ+ group exhibits persistently higher subjective than objective memory performance, resulting in positive discrepancy scores indicative of overestimation. Standardized scores for the Objective and Subjective metric are based on the reference group at individual baseline. The Discrepancy metric (i.e., Subjective–Objective) is not further standardized. Note that the Weschler scales and ECOG availability varied across ADNI visits/phases.

**Table 2.** Individual Baseline Performance on the Different Cognitive Tests in the Aβ+, Aβ− and Reference Groups.

| **Group** | **Sex** | ***n*** | **ECOG** | **Subjective Z-score** | **Weschler Immediate / Z-score** | **Weschler Delayed / Z-score** | **Weschler average** | **Discrepancy** |
|---|---|---|---|---|---|---|---|---|
| **Aβ+** | Female | 154 | 17.77 (5.96) | -0.79 (1.3) | 9.45 (5.49) / -1.65 (1.56) | 6.44 (6.3) / -1.89 (1.59) | -1.77 (1.53) | 1.12 (1.84) |
|  | Male | 196 | 19.07 (5.83) | -1.07 (1.27) | 8.57 (4.69) / -1.9 (1.33) | 5.85 (5.17) / -2.04 (1.3) | -1.97 (1.27) | 1.08 (1.61) |
| **Aβ-** | Female | 147 | 15.8 (5.12) | -0.36 (1.12) | 14.01 (4.22) / -0.36 (1.2) | 12.45 (4.69) / -0.38 (1.18) | -0.37 (1.16) | -0.02 (1.45) |
|  | Male | 163 | 16.24 (5.24) | -0.45 (1.14) | 12.67 (4.34) / -0.74 (1.23) | 11.01 (5.04) / -0.75 (1.27) | -0.74 (1.22) | 0.36 (1.49) |
| **Reference** | Female | 91 | 14.48 (4.72) | -0.05 (1.00) | 15.77 (3.59) / 0.10 (1.03) | 14.75 (3.98) / 0.12 (1.02) | 0.11 (0.99) | -0.16 (1.30) |
|  | Male | 89 | 14.02 (4.70) | 0.05 (1.00) | 15.08 (3.38) / -0.10 (0.97) | 13.82 (3.78) / -0.12 (0.97) | -0.11 (0.94) | 0.16 (1.39) |

*Note.* ECOG = Everyday Cognition Scale. *Z*-scoring based on the reference sample. N = number of participants. The reference group is the cognitively unimpaired Aβ− group.

### Brain Volume Trajectories Across Groups (eq. 2)

#### Precuneus

The Aβ+ group exhibited lower precuneus volume compared with the Aβ− group (*β* = −0.592, *p* < 0.001). There were no sex differences in precuneus volume (*β* = 0.174, *p* = 0.282). There was a significant Aβ status × Time interaction (*β* = −0.216, *p* < 0.001), indicating greater precuneus volume decline over time in the Aβ+ group. Precuneus volume also declined significantly over time in the Aβ− group (*β* = −0.137, *p* < 0.001).

#### Control regions

No significant difference in PCC volume was observed between groups (*β* = −0.085, *p* = 0.188). There were no sex differences PCC volume (*β* = 0.028, *p* = 0.696). There was a small but significant Aβ status × Time interaction (*β* = −0.094, *p* < 0.001), indicating greater PCC volume decline in the Aβ+ group compared with the Aβ− group. PCC volume showed a small but significant increase over time in the Aβ− group (*β* = 0.038, *p* < 0.001).

The Aβ+ group showed significantly lower HC volume compared with the Aβ− group (*β* = −0.686, *p* < 0.001). There were no sex differences in HC volume (*β* = −0.203, *p* = 0.003). A significant Aβ status × Time interaction was also observed (*β* = −0.226, *p* < 0.001), indicating greater HC volume decline over time in the Aβ+ group. HC volume also declined significantly over time in the Aβ− group (*β* = −0.186, *p* < 0.001).

The Aβ+ group exhibited lower SFG volume compared with the Aβ− group (*β* = −0.747, *p* < 0.001). Females had larger SFG volumes than males overall (*β* = −0.508, *p* = 0.032). The Aβ status × Time interaction was significant (*β* = −0.110, *p* < 0.01), demonstrating greater SFG volume decline in the Aβ+ group. SFG volume also declined significantly over time in the Aβ− group (*β* = −0.192, *p* < 0.001). Table 3 shows the regional brain volumes in both groups and in the reference sample at baseline, stratified by Sex.

**Table 3.** Brain Volume Measurements at Baseline for Both Amyloid Groups and the Reference Sample.

| **Group** | **Sex** | ***n*** | **Precuneus** | **Hippocampus** | **SFG** | **PCC** | **ICV** |
|---|---|---|---|---|---|---|---|
| Aβ+ | Female | 154 | 21.77 (2.32) | 6.21 (0.81) | 27.46 (3.44) | 9.49 (1.07) | 1324.44 (99.03) |
|  | Male | 196 | 25.1 (2.92) | 6.6 (0.93) | 30.92 (3.97) | 10.95 (1.27) | 1505.92 (117.73) |
| Aβ− | Female | 153 | 22.6 (2.39) | 6.73 (0.85) | 28.65 (3.17) | 9.73 (1.03) | 1346.54 (103.97) |
|  | Male | 175 | 25.19 (3.04) | 7.06 (0.95) | 31.19 (3.57) | 10.75 (1.24) | 1488.56 (114.4) |
| Reference | Female | 91 | 22.24 (2.48) | 6.78 (0.71) | 28.15 (3.42) | 9.72 (1.01) | 1338.09 (103.25) |
|  | Male | 89 | 25.53 (3.15) | 7.27 (0.90) | 31.07 (3.92) | 10.74 (1.24) | 1493.25 (125.53) |

*Note.* All regional brain volume values are in cm<sup>3</sup>. SFG = Superior frontal gyrus. PCC = Posterior cingulate cortex. ICV = Intracranial volume. N = number of participants. The reference group is the cognitively unimpaired Aβ− group.

### Effect of Sex on the Relationship Between Precuneus Volume and Memory Discrepancy in Aβ− and Aβ+ Groups

**Cross-sectional analyses (eq. 3)** examined whether the association between precuneus volume and subjective–objective memory discrepancy differed by sex at baseline (Table 4). In the Aβ+ group, the Precuneus volume × Sex interaction was marginally significant (*β* = 0.128, *p* = 0.065). Precuneus volume was significantly associated with Discrepancy (*β* = −0.220, *p* < 0.001). Among covariates, higher baseline CDR−SB scores were associated with greater Discrepancy (*β* = 0.282, *p* < 0.001), but there was no main effect of Sex (*β* = 0.401, *p* = 0.086). The model explained 12.6% of the variance. In the Aβ− group, no significant Baseline precuneus volume × Sex interaction was observed (*β* = −0.088, *p* = 0.148), and Precuneus volume was not significantly associated with Discrepancy (*β* = 0.086, *p* = 0.144). In contrast, there was a main effect of Sex on Discrepancy (*β* = 0.527, *p* = 0.007), and baseline CDR−SB was also significantly associated with Discrepancy (*β* = 0.242, *p* < 0.002). The overall model explained 10.1% of the variance. Table 4 shows the linear regression estimates for both amyloid groups.

**Table 4.** Linear Regression Estimates for the cross-sectional model (eq. 3) Examining the Relationships Between Precuneus Volume and Sex on Memory Discrepancy, Stratified by Amyloid Status.

| **Aβ+** |  |  |  |  |
|---|---|---|---|---|
| ***term*** | ***Estimate*** | ***SE*** | ***t*** | ***p*** |
| (Intercept) | -0.501 | 1.587 | -0.316 | 0.752 |
| Precuneus volume | -0.220 | 0.065 | -3.374 | **< 0.001** |
| Sex | 0.401 | 0.233 | 1.724 | 0.086 |
| CDRSB | 0.282 | 0.054 | 5.224 | **< 0.001** |
| Education | 0.003 | 0.034 | 0.100 | 0.921 |
| ICV | 0.000 | 0.001 | 0.445 | 0.657 |
| Precuneus volume: Sex | 0.128 | 0.069 | 1.852 | 0.065 |
| **Aβ−** |  |  |  |  |
| ***term*** | **Estimate** | **SE** | ***t*** | ***p*** |
| (Intercept) | 4.337 | 1.519 | 2.855 | **0.005** |
| Precuneus volume | 0.086 | 0.059 | 1.463 | 0.144 |
| Sex | 0.527 | 0.193 | 2.734 | **0.007** |
| CDRSB | 0.242 | 0.076 | 3.169 | **0.002** |
| Education | -0.136 | 0.033 | -4.127 | **< 0.001** |
| ICV | -0.002 | 0.001 | -1.508 | 0.133 |
| Precuneus volume: Sex | -0.088 | 0.061 | -1.451 | 0.148 |

*Note.* CDR−SB: Clinical Dementia Rating – Sum of Boxes. ICV: Intracranial Volume. Significant results in bold.

**Longitudinal models (eq. 4).** In the Aβ+ group, the main effect of within-person Precuneus delta volume was significant (*β* = −0.541, *p* < 0.001), indicating that a decrease in precuneus volume over time was associated with increased Discrepancy scores in females (the reference group). See Table 5. The interaction between Precuneus delta volume and Sex was also significant (*β* = 0.508, *p* = 0.002), indicating that this association was attenuated in males compared to females. Baseline precuneus volume was also significantly associated with Discrepancy (*β* = −0.146, *p* < 0.001), suggesting that lower Baseline precuneus volume was associated with greater Discrepancy. Sex was also a significant predictor of Discrepancy (*β* = 0.448, *p* = 0.035), indicating greater Discrepancy in males. The marginal R^2^ was 0.125, while the conditional R^2^ was 0.769.

**Table 5.** Longitudinal Precuneus Model Results for Both Amyloid Groups.

| **Aβ+** |  |  |  |  |  |
|---|---|---|---|---|---|
| ***term*** | ***Estimate*** | ***SE*** | ***t*** | ***df*** | ***p*** |
| (Intercept) | 2.688 | 1.014 | 2.651 | 355.448 | **0.008** |
| Precuneus (baseline) | -0.146 | 0.041 | -3.532 | 355.465 | **< 0.001** |
| Precuneus_delta | -0.540 | 0.135 | -4.007 | 743.741 | **< 0.001** |
| Sex | 0.448 | 0.211 | 2.118 | 378.863 | **0.035** |
| CDRSB_baseline | 0.333 | 0.051 | 6.576 | 355.029 | **< 0.001** |
| Time | 0.215 | 0.085 | 2.540 | 154.153 | **0.012** |
| Education | -0.001 | 0.031 | -0.043 | 342.870 | 0.966 |
| ICV | 0.001 | 0.001 | 0.720 | 369.819 | 0.472 |
| Precuneus_delta: Sex | 0.508 | 0.167 | 3.042 | 667.881 | **0.002** |
| **Aβ-** |  |  |  |  |  |
| **term** | **Estimate** | ***SE*** | ***t*** | ***df*** | ***p*** |
| *(Intercept*) | 2.637 | 0.953 | 2.769 | 329.163 | **0.006** |
| Precuneus (baseline) | -0.004 | 0.038 | -0.105 | 322.947 | 0.917 |
| Precuneus_delta | 0.031 | 0.116 | 0.271 | 900.690 | 0.786 |
| Sex | 0.558 | 0.178 | 3.144 | 324.161 | **0.002** |
| CDRSB_baseline | 0.242 | 0.074 | 3.292 | 334.928 | **0.001** |
| Time | -0.006 | 0.045 | -0.133 | 155.442 | 0.894 |
| Education | -0.131 | 0.030 | -4.331 | 307.671 | **< 0.001** |
| ICV | -0.001 | 0.001 | -0.553 | 337.229 | 0.581 |
| Precuneus_delta: Sex | 0.123 | 0.152 | 0.811 | 851.986 | 0.418 |

*Note.* Marginal *R*<sup>2</sup> = 0.125. Conditional *R*<sup>2</sup> = 0.769. CDR−SB: Clinical Dementia Rating – Sum of Boxes. ICV: Intracranial Volume. Significant values in bold.

In the Aβ− group, within-person precuneus delta volume was not associated with Discrepancy (*β* = 0.031, *p* = 0.786), nor was Baseline precuneus volume (*β* = −0.004, *p* = 0.917). Sex was the only significant predictor of Discrepancy (*β* = 0.558, *p* = 0.002), indicating greater Discrepancy in males. The interaction between within-person Precuneus volume change and Sex was not significant (*β* = 0.123, *p* = 0.418), indicating that the association between longitudinal precuneus volume change and Discrepancy did not differ by Sex. The marginal R^2^ was 0.087, while the conditional R^2^ was 0.714. Table 5 shows the results of the longitudinal precuneus models. Figure 3 shows the interaction results for both amyloid groups.

**Figure 3.**
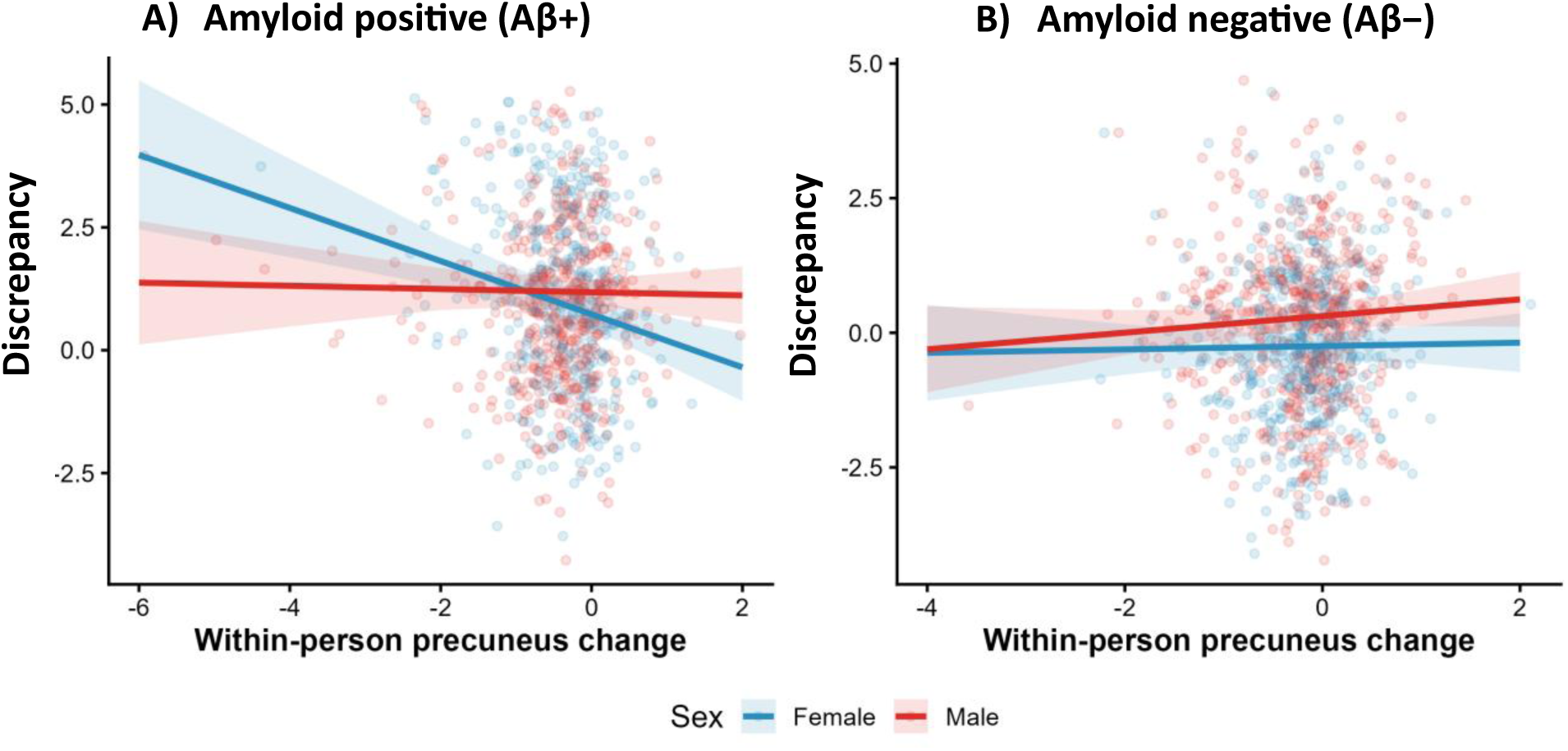
Relationship between Within-person precuneus volume change (in cm^3^) and Discrepancy, stratified by Sex and by amyloid status. Note. Sex-specific association between longitudinal precuneus change and discrepancy scores according to amyloid status. Adjusted model predictions from linear mixed-effects models are shown as lines with 95% confidence intervals, with raw observations overlaid. Separate panels depict amyloid-positive (Aβ+) in A) and amyloid-negative (Aβ−) in B). For the Aβ+ group, influence diagnostics were performed using Cook’s distance at the participant level. No highly influential observations were identified (maximum Cook’s distance = 0.028). A sensitivity analysis excluding participants exceeding the 4/n threshold yielded comparable results, with the Precuneus (atrophy) × Sex interaction remaining significant (*β* = 0.680, *p* < 0.001).

### Control Analyses

Our findings suggest that greater precuneus delta (i.e., atrophy over time) is associated with greater memory discrepancy and that this relationship is stronger in females. To determine if this sex difference is unique to the effect of atrophy of the precuneus, we conducted a series of control analyses using other brain regions known to play a role in memory.

#### Posterior Cingulate Cortex

In the Aβ+ group, longitudinal PCC delta was significantly associated with longitudinal changes in Discrepancy after adjustment for baseline PCC volume, CDR-SB, Time, Education, and ICV (*β* = −1.008, *p* = 0.001). Baseline PCC volume was also significantly associated with Discrepancy (*β* = −0.263, *p* = 0.006), whereas Sex was not a significant predictor of Discrepancy (*β* = 0.307, *p* = 0.142). The PCC atrophy × Sex interaction was not significant (*β* = 0.622, *p* = 0.105), indicating no evidence that the association between longitudinal PCC decline and Discrepancy differed by Sex.

In the Aβ− group, neither longitudinal PCC delta (*β* = 0.236, *p* = 0.346) nor baseline PCC volume (*β* = 0.092, *p* = 0.287) was associated with Discrepancy. Sex was a significant predictor of Discrepancy in this model (*β* = 0.518, *p* = 0.003). The PCC delta × Sex interaction was not significant (*β* = −0.068, *p* = 0.836), indicating no evidence that the association between longitudinal PCC delta and Discrepancy differed by Sex.

#### Hippocampus

In the Aβ+ group, lower baseline HC volume was associated with greater Discrepancy (*β* = −0.340, *p* = 0.001), and greater longitudinal HC delta was also associated with greater Discrepancy (*β* = −0.739, *p* < 0.001). Sex was not a significant predictor of Discrepancy in this model (*β* = 0.251, *p* = 0.26). The HC delta × Sex interaction was not significant (*β* = 0.142, *p* = 0.581), indicating that the association between longitudinal HC decline and Discrepancy did not differ by Sex.

In the Aβ− group, lower baseline HC volume (*β* = −0.274, *p* = 0.002) and greater longitudinal HC delta (*β* = −0.645, *p* = 0.012) were both associated with greater Discrepancy. Sex was also a significant predictor of Discrepancy (*β* = 0.455, *p* = 0.013). The HC delta × Sex interaction was not significant (*β* = −0.059, *p* = 0.841), indicating that the association between longitudinal HC decline and Discrepancy did not differ by Sex. Together, these findings suggest that both baseline and longitudinal HC measures were associated with Discrepancy in both amyloid groups, with no evidence that the associations between HC measures and Discrepancy differed by Sex.

#### Superior Frontal Gyrus

In the Aβ+ group, greater longitudinal SFG delta was associated with greater change in Discrepancy scores after adjustment for baseline SFG volume, CDR-SB, Time, Education, and ICV (*β* = −0.194, *p* = 0.008). Baseline SFG volume was not significantly associated with Discrepancy (*β* = −0.011, *p* = 0.698), and Sex was not a significant predictor of Discrepancy (*β* = 0.281, *p* = 0.185). The SFG delta × Sex interaction reached marginal significance (β = 0.182, p = 0.079), suggesting no clear statistically significant evidence that the association between SFG decline and Discrepancy differed by Sex.

In the Aβ− group, neither longitudinal SFG change (*β* = 0.039, *p* = 0.617) nor baseline SFG volume (*β* = −0.035, *p* = 0.200) was associated with Discrepancy. Sex was a significant predictor of Discrepancy (*β* = 0.491, *p* = 0.006). The SFG delta × Sex interaction was not significant (*β* = −0.121, *p* = 0.231), indicating no evidence that the association between longitudinal SFG change and Discrepancy differed by Sex.

## Discussion

This analysis aimed to investigate the potential neural substrates underlying sex differences in the discrepancy between subjective and objective memory performance observed in AD. We conducted an analysis of ADNI data to compare the relationship between precuneus volume and memory discrepancy in males and females and stratified our analyses by amyloid status. We hypothesized that the precuneus, given its established role in metamemory processes and evidence of early sex-related vulnerability in AD, would show a preferential association with discrepancy measures compared with other regions implicated in either AD progression or meta-memory abilities, namely the HC, the PCC, and the SFG. Specifically, we hypothesized that greater preservation of precuneus integrity would be associated with a smaller subjective–objective memory discrepancy in females, reflecting a potential protective effect of preserved precuneus structure against increasing discrepancy.

Functional neuroimaging has long established the precuneus as a central hub of the DMN, serving essential roles in self-referential processing, and memory (Dadario & Sughrue, 2023). The female-specific association between precuneus atrophy and increasing cognitive discrepancy may reflect sexually dimorphic patterns of DMN engagement during memory processes. Recent evidence suggests that memory retrieval relies more on intra-DMN circuitry in women, with the PCC/precuneus emerging as the strongest predictor amongst other targeted regions, while males show greater reliance on visual network circuitry (Ju et al., 2023). This greater reliance on the precuneus integrity for females may be particularly important for memory-related self-monitoring, such that progressive structural disruption within this network could disproportionately affect the alignment between subjective and objective memory performance in this group.

This potential sex-specific vulnerability may be particularly relevant in the context of Aβ pathology, given evidence that amyloid and APOE ε4 status interact with sex to shape DMN connectivity, with effects emerging in females but not males (Caldwell et al., 2019). Notably, early Aβ accumulation preferentially disrupts DMN regions such as the precuneus, which supports metacognitive monitoring (Palmqvist et al., 2017; Ye et al., 2018). Our finding that lower precuneus volume was associated with greater cognitive discrepancy specifically among Aβ+ females fits this pattern, suggesting that preserved precuneus integrity may be especially important for maintaining alignment between subjective and objective memory performance in females with AD pathology. The absence of precuneus-related effects among Aβ− individuals further suggests that preserved precuneus volume alone may be insufficient to influence discrepancy in the absence of AD-related pathology. Instead, Aβ pathology may unmask the functional relevance of the precuneus by disrupting DMN connectivity, consistent with evidence that early Aβ-related alterations in the precuneus and other DMN hubs reflect a breakdown of compensatory network organization (Elman et al., 2016). Further support for the sensitivity of the precuneus to Aβ-related disruption comes from evidence of abnormally elevated task-related activation during memory retrieval, with greater precuneus activity associated with higher Aβ burden in cognitively normal APOE4 carriers (Fischer et al., 2025).

The control-region pattern is broadly consistent with known patterns of amyloid progression. HC volume was linked to discrepancy in both Aβ+ and Aβ− groups, consistent with evidence that hippocampal tau, which precedes HC atrophy, is comparatively decoupled from global amyloid burden (Aumont et al., 2025). This may indicate a more domain-general memory-encoding contribution rather than an amyloid-specific mechanism. The PCC, by contrast, was linked to discrepancy only in the Aβ+ group, with no difference between sexes, matching PET-staging evidence that the posterior cingulate/precuneus are among the earliest and most consistently amyloid-affected regions (Pfeil et al., 2021), and likely reflects shared vulnerability with the precuneus given their tight coupling. Finally, the SFG showed only a weak, longitudinal-only effect in Aβ+ that did not depend on sex, consistent with lateral frontal cortex being reached later in amyloid progression, after posterior DMN hubs are already affected (Chung et al., 2022).

This staged pattern situates the precuneus within a coherent account: HC’s link is largely amyloid-independent, SFG’s would emerge only later and weakly, and PCC and precuneus (the earliest amyloid targets) are the only regions showing amyloid-dependent discrepancy associations. Of these two, only the precuneus is sex-dependent, indicating that early posterior amyloid burden is necessary but not sufficient for the sex effect. Something specific to the precuneus, plausibly because of its sex-dimorphic role in DMN-based memory retrieval, may drive the interaction (Ju et al., 2023).

Our results demonstrate that progressive increases in cognitive discrepancy were specific to the Aβ+ group, whereas Aβ− individuals maintained relatively stable alignment between subjective and objective memory performance during follow-up. This suggests that Aβ pathology represents a critical context in which self-monitoring of cognitive decline becomes progressively disrupted. Although longitudinal atrophy across all four targeted regions were associated with increased discrepancy in Aβ+ individuals, only the precuneus demonstrated a significant sex-dependent relationship. Specifically, greater longitudinal precuneus atrophy was associated with increasing discrepancy in females, whereas this association was attenuated in males. Although the precuneus atrophy-by-sex interaction was also observed for objective memory performance, the interaction remained significant for discrepancy after adjusting for objective memory, indicating that the precuneus contributes to variance in subjective– objective mismatch beyond its relationship with objective memory impairment alone. Taken together, these findings reveal that the precuneus structural integrity acts as a sex-specific marker for metacognitive monitoring in the context of AD pathology.

### Limitations

An important limitation in this study is about additional brain areas that could have been targeted.

The insula, other cingulate areas, other regions of the DMN or even frontal areas that are all linked to metacognitive processes (Hallam et al., 2020) could have been interesting candidates in their potential role in anosognosia in AD. Another limitation is that our analysis only looked at structural integrity, inferred from atrophy and volume measurements. Assessing functional integrity may have provided additional and complementary insights, as previous studies on anosognosia in AD have demonstrated, particularly within DMN regions (Therriault et al., 2018).

### Future Directions

It is, first, crucial to reproduce and replicate these findings in other similar datasets such has PreventAD, and CIMAQ (Belleville et al., 2019; Villeneuve et al., 2025). While the precuneus was investigated, whole-brain structural analyses (e.g., voxel-wise morphometry) could provide a better answer to regions responsible for sex-differential discrepancy in AD. Potentially, a metamemory network in the style of the DMN, but involved in metacognition and its insight could be elucidated.

## Conclusion

This study demonstrates that sex and Aβ pathology shape the relationship between brain structure and subjective–objective memory discrepancy in AD. The precuneus showed a unique sex- and Aβ- dependent association with discrepancy, particularly in females with Aβ pathology. These findings suggest that preserved precuneus integrity may be particularly relevant to metacognitive monitoring in females with AD pathology and highlight the importance of considering sex and pathology when studying cognitive insight in AD.

## Acknowledgments, others

The author would like to acknowledge its source of funding: the Alzheimer’s Society of Canada for its doctoral scholarship to first author MAAO, and La famille Louise et André Charron for their funding in the curation and preparation of this article.

Data collection and sharing for this project was funded by the Alzheimer’s Disease Neuroimaging Initiative (ADNI) (National Institutes of Health Grant U01 AG024904) and DOD ADNI (Department of Defense award number W81XWH-12-2-0012). ADNI is funded by the National Institute on Aging, the National Institute of Biomedical Imaging and Bioengineering, and through generous contributions from the following: AbbVie, Alzheimer’s Association; Alzheimer’s Drug Discovery Foundation; Araclon Biotech; BioClinica, Inc.; Biogen; Bristol-Myers Squibb Company; CereSpir, Inc.; Cogstate; Eisai Inc.; Elan Pharmaceuticals, Inc.; Eli Lilly and Company; EuroImmun; F. Hoffmann-La Roche Ltd and its affiliated company Genentech, Inc.; Fujirebio; GE Healthcare; IXICO Ltd.; Janssen Alzheimer Immunotherapy Research & Development, LLC.; Johnson & Johnson Pharmaceutical Research & Development LLC.; Lumosity; Lundbeck; Merck & Co., Inc.; Meso Scale Diagnostics, LLC.; NeuroRx Research; Neurotrack Technologies; Novartis Pharmaceuticals Corporation; Pfizer Inc.; Piramal Imaging; Servier; Takeda Pharmaceutical Company; and Transition Therapeutics. The Canadian Institutes of Health Research is providing funds to support ADNI clinical sites in Canada. Private sector contributions are facilitated by the Foundation for the National Institutes of Health (www.fnih.org). The grantee organization is the Northern California Institute for Research and Education, and the study is coordinated by the Alzheimer’s Therapeutic Research Institute at the University of Southern California. ADNI data are disseminated by the Laboratory for Neuro Imaging at the University of Southern California.

## Author Note

Data curation and analysis were supported by a CIHR Project Grant FRN 165921 and a donation from La Famille Louise et Andre Charron. A portion of these findings have been presented at the Alzheimer’s Association International Conference in July 2024. We have no conflicts of interest to disclose.

